# The Personal Genome Project-UK: an open access resource of human multi-omics data

**DOI:** 10.1101/566711

**Authors:** Olga Chervova, Lucia Conde, José Afonso Guerra-Assunção, Ismail Moghul, Amy P. Webster, Alison Berner, Elizabeth Larose Cadieux, Yuan Tian, Vitaly Voloshin, Rifat Hamoudi, Javier Herrero, Stephan Beck

## Abstract

Integrative analysis of multi-omics data is a powerful approach for gaining functional insights into biological and medical processes. Conducting these multifaceted analyses on human samples is often complicated by the fact that the raw sequencing output is rarely available under open access. The Personal Genome Project UK (PGP-UK) is one of few resources that recruits its participants under open consent and makes the resulting multi-omics data freely and openly available. As part of this resource, we describe the PGP-UK multi-omics reference panel consisting of ten genomic, methylomic and transcriptomic data. Specifically, we outline the data processing, quality control and validation procedures which were implemented to ensure data integrity and exclude sample mix-ups. In addition, we provide a REST API to facilitate the download of the entire PGP-UK dataset. The data are also available from two cloud-based environments, providing platforms for free integrated analysis. In conclusion, the genotype-validated PGP-UK multi-omics human reference panel described here provides a valuable new open access resource for integrated analyses in support of personal and medical genomics.

## Background & Summary

The Personal Genome Project UK (PGP-UK) is a member of the global PGP network together with the PGPs in the United States, Canada, Austria and China. The PGP network aims to provide multi-omics and trait data under open access to the community. This contributes to personalised medicine by advancing our understanding of how phenotypes and the development of diseases are influenced by genetic, epigenetic, environmental and lifestyle factors. While all five PGP centres generate whole-genome sequencing (WGS), some PGPs, such as PGP-UK, produce additional multi-omics data.

To participate in this study, volunteers must pass the eligibility criteria (e.g. be a UK citizen or permanent resident), sign the open consent form and pass a very thorough entrance exam. The objective of the exam is to ensure that the participant understands the key PGP-UK procedures and the potential risks of being involved in a project of this nature. At present, 1100 subjects have successfully enrolled in the project, and over a hundred of them have had their genomes sequenced. Once enrolled, participants are invited for sample collection which involves giving a blood or saliva sample or both for DNA and RNA extraction. DNA sequencing is then performed followed by data analysis. The results are reported back to the participants in the form of a Genome Report that is made publicly available after a grace period of one month. However, the participant is able to withdraw from the project at any time. DNA methylation data is generated using the Illumina HumanMethylation450 array (450k) and results are displayed in a freely available Methylome Report, a unique feature of the UK branch of the project. The preparation of both Genome and Methylome reports is discussed in more details in the Usage Notes Section.

A pilot cohort of ten members of the public make up the PGP-UK multiomics reference panel. For this cohort, we collected whole-genome bisulfite sequencing (WGBS) and RNA sequencing (RNA-seq) in addition to WGS and 450k data. Figure 1 shows a schematic of the PGP-UK workflow. More information about PGP-UK can be found in [18, 5] and on the project’s website www.personalgenomes.org.uk.

**Figure 1:**
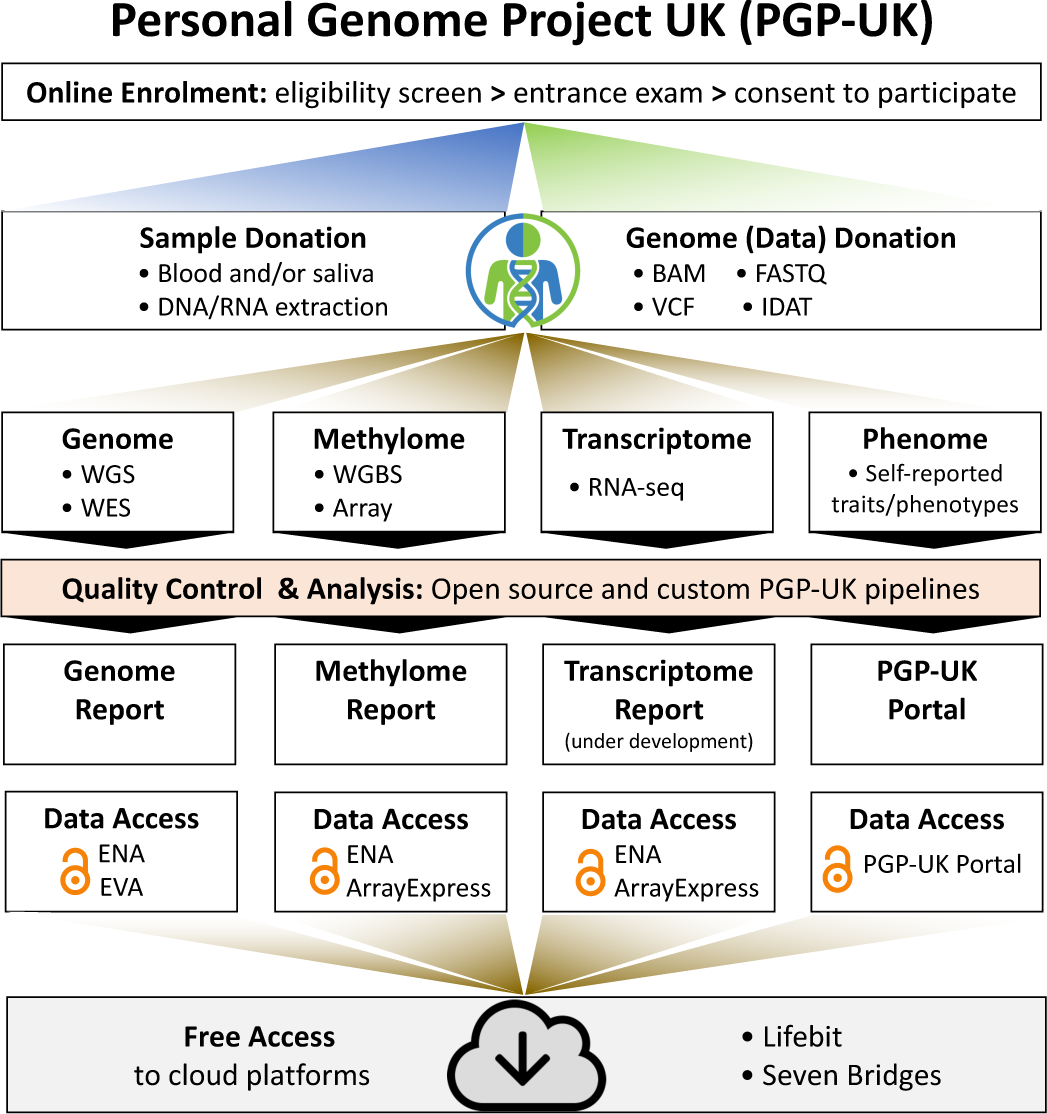
PGP-UK workflow

While controlled access multi-omics data can be submitted into a single public repository (e.g. EGA in Europe or dbGaP in the USA), there is currently no single public repository for open access multi-omics data. Consequently, the different types of datasets (WGS, WGBS, RNA-seq, 450k) were submitted to the corresponding repositories (ENA, EVA, ArrayExpress) at EMBL-EBI. The details are given in the Data Records section and direct data download links are provided on the PGP-UK data web page www.personalgenomes.org.uk/data. For convenience, we offer a web API to download all the available PGP-UK data (see Data Records). The cumulative size of the PGP-UK multi-omics reference panel exceeds 2TB, which means that it would take over 10 days to download (with mean UK download speed of 18.57Mbps, Ofcom 2018). To overcome this limitation, we collaborated with two cloud platform providers (Seven Bridges Genomics and Lifebit) to host PGP-UK data in their respective clouds for unrestricted access as briefly described in the Data Records section.

In this paper, we describe the PGP-UK multi-omics human reference panel derived from 10 participants. We followed best practices to perform various quality control (QC) checks to ensure the quality of the pilot WGS, WGBS, RNA-seq and 450k datasets as described in the Technical Validation section. Finally, we describe the methods employed for multi-omics data matching, which ensure that samples are mapped to the correct participant.

## Methods

### Ethics

The PGP-UK study is approved by the University College London (UCL) Research Ethics Committee (ID Number 4700/001) subject to annual reviews and renewals. All the research activities in the project are conducted in accordance with the Declaration of Helsinki, UK national laws and medical research regulatory requirements. Prior to their enrolment, every participant must pass an entrance exam, give their consent to participate in the project and agree for their data and associated reports to be made publicly available under open access.

### Tissue Samples

Blood samples were collected using EDTA Vacutainers (Becton Dickinson). Saliva samples were collected using Oragene OG-500 self-sampling kits. Sample processing and storage protocols were in line with HTA-approved standard operating procedures.

### Whole-genome sequencing (WGS)

WGS libraries were prepared from whole blood DNA using Illumina TruSeq Nano in accordance with standard operating procedures. Sequencing was performed on an Illumina HiSeq X Ten platform with an average depth of 30X. The resulting reads were trimmed using TrimGalore software, mapped to the human reference genome hg19 (GRCh37) using BWA-MEM algorithm (BWA v.0.7.12 [13]). Ambiguously mapped reads (MAPQ<10) and duplicated reads were removed using `SAMtools` v.1.2 [14] and `Picard` v.1.130 respectively. Genomic variants were called following the `Genome Analysis Toolkit` software (`GATK` v.3.4-46) best practices.

The corresponding FASTQ, BAM and VCF files were deposited in European Nucleotide Archive (ENA), see Data Citation 1.

### Whole-genome bisulfite sequencing (WGBS)

DNA was extracted from blood samples followed by bisulfite conversion and library preparation using the TruMethyl Whole Genome Kit v2.1. WGBS was performed on an Illumina HiSeq X Ten platform with an average depth of 15X. Generated FASTQ files were processed using GemBS v.0.11.7 software [15].

Resulting FASTQ and BAM files were deposited in the European Nucleotide Archive (ENA), see Data Citation 1.

### RNA Sequencing (RNA-seq)

RNA-seq was performed using 20ng of RNA isolated from whole blood. All the involved procedures were implemented in accordance with the corresponding manufacturers’ protocols.

Libraries for RNA-seq were prepared with SENSE mRNA-seq Library Prep Kit v2, purified and amplified (18 PCR cycles). After adding adapters and indices, sequencing libraries were further purified using Solid Phase Reversible Immobilisation beads. The output was QC-verified and quantified using Qubit fluorometer. Finally, libraries were QC-analysed on Bioanalyzer 2100 and further quantified by qPCR with KAPA library quantification kit and the sequencing was performed on Illumina HiSeq 4000.

RNA-seq FASTQ files are available to download from the ArrayExpress and European Nucleotide Archive repositories, see Data Citations 2 and 3.

### DNA Methylation Profiling

Genomic DNA (500ng) extracted from whole blood and saliva was bisulfite converted using the EZ DNA Methylation Kit (Zymo Research) following the recommended incubation conditions for Illumina Infinium HumanMethylation450 BeadChip (450k). Methylation profiling was subsequently performed on 450k arrays using Illumina iScan Microarray Scanner at UCL Genomics, in accordance with standard operating procedures.

Raw DNA methylation array data (IDAT files) for PGP-UK participants were submitted to the ArrayExpress repository, see Data Citation 4.

## Data Records

The entire PGP-UK dataset is freely available for download from public repositories with no access restrictions (see Data Citations). Links for the particular datasets are provided on the PGP-UK website (www.personalgenomes.org.uk). Accession numbers and dataset identifiers are presented in Table 1. Basic phenotype data, which includes self-reported age, sex, smoking status, etc., alongside with genome and methylome reports, generated by the PGP-UK, can be found on the project’s data web page www.personalgenomes.org.uk/data. Further-more, all of the data (including associated metadata) are available through the PGP-UK API. The API is compliant with the Open API Specification 3.0 and is documented at www.personalgenomes.org/api.

**Table 1:** PGP-UK data identifiers. The table contains ENA accession numbers for WGS, WGBS and RNA-seq, for 450k data it shows Sentrix IDs and positions.

| Sample ID | EBI ID | Tissue | WGS | WGBS | 450k | RNA-seq |
| --- | --- | --- | --- | --- | --- | --- |
|  |  |  | ENA<br>PRJEB17529 | ENA<br>PRJEB17529 | ArrayExpress<br>E-MTAB-5377 | ENA<br>PRJEB25139 |
| uk35C650 | SAMEA4545245 | blood | ERX1796409 | ERX2408504 | 101130760050_R04C02 | ERX2373318 |
|  |  | saliva |  |  | 101130760049_R03C01 |  |
| uk2E2AAE | SAMEA4545246 | blood | ERX1796410 | ERX2408505 | 101130760050_R05C02 | ERX2373321 |
|  |  | saliva |  |  | 101130760050_R03C01 |  |
| uk2DF242 | SAMEA4545247 | blood | ERX1796411 | ERX2408506 | 101130760049_R06C02 | ERX2373317 |
|  |  | saliva |  |  | 101130760049_R03C02 |  |
| uk740176 | SAMEA4545248 | blood | ERX1796412 | ERX2408507 | 101130760050_R06C02 | ERX2373324 |
|  |  | saliva |  |  | 101130760050_R06C01 |  |
| uk33D02F | SAMEA4545249 | blood | ERX1796413 | ERX2408508 | 101130760049_R05C02 | ERX2373316 |
|  |  | saliva |  |  | 101130760049_R04C02 |  |
| uk0C72FF | SAMEA4545250 | blood | ERX1796414 | ERX2408509 | 101130760049_R06C01 | ERX2373322 |
|  |  | saliva |  |  | 101130760050_R01C01 |  |
| uk1097F9 | SAMEA4545251 | blood | ERX1796415 | ERX2408510 | 101130760050_R02C01 | ERX2373320 |
|  |  | saliva |  |  | 101130760050_R01C02 |  |
| uk174659 | SAMEA4545252 | blood | ERX1796416 | ERX2408511 | 101130760050_R05C01 | ERX2373325 |
|  |  | saliva |  |  | 101130760049_R05C01 |  |
| uk85AA3B | SAMEA4545253 | blood | ERX1796417 | ERX2408512 | 101130760049_R02C02 | ERX2373323 |
|  |  | saliva |  |  | 101130760049_R01C01 |  |
| uk481F67 | SAMEA4545254 | blood | ERX1796418 | ERX2408513 | 101130760049_R02C01 | ERX2373319 |
|  |  | saliva |  |  | 101130760050_R02C02 |  |

Whole genome sequencing and whole genome bisulfite sequencing data are freely available from the European Nucleotide Archive (ENA) under the project ID PRJEB17529 (Dataset Citation 1). RNA-seq data is deposited in ArrayExpress under the accession number E-MTAB-6523 (Data Citation 2) and in ENA PRJEB25139 (Data Citation 3). DNA methylation array data for PGP-UK participants is stored in ArrayExpress under the accession number E-MTAB-5377 (Data Citation 4).

The PGP-UK pilot dataset described in [18] resulted in the PGP-UK multi-omics reference panel described here. The datasets are available from the above-mentioned repositories and from the Seven Bridges Cancer Genomics cloud (docs.cancergenomicscloud.org/docs/personal-genome-project-uk-pgp-uk-pilot-dataset), which offers various tools and workflows for genomic and epigenomic data analysis.

The PGP-UK multi-omics reference panel is also available in the Lifebit cloud through their Open Data project (opendata.lifebit.ai/table/pgp) along with interactive analyses (ancestry, phenotypic traits, genetic variance) and custom pipelines provided by Lifebit’s cloud-computing platform Deploit (`deploit.lifebit.ai`).

## Technical Validation

In this section, we describe the outcomes of the PGP-UK data quality control checks and validation for the PGP-UK pilot cohort. In a first instance, we describe the QC framework and discuss outputs for each types of data collected. Then, we provide details of multi-omics data matching validation procedures based on cross-comparison of variants between different data types for each individual.

### Data Quality Control

#### WGS data QC

Quality control of the reported WGS data was performed using `FastQC` v.0.11.2 and `Picard` v.1.130 tools. QC reports were generated using `MultiQC` v.1.5 software [8].

WGS average median coverage is above 35X (varies between 30X and 47X across samples) with more than 73% of the bases covered reaching 30X or more (varies between 54% and 95% across samples), see Figure 2a. A summary of the WGS QC analysis is presented in Table 2.

**Figure 2:**
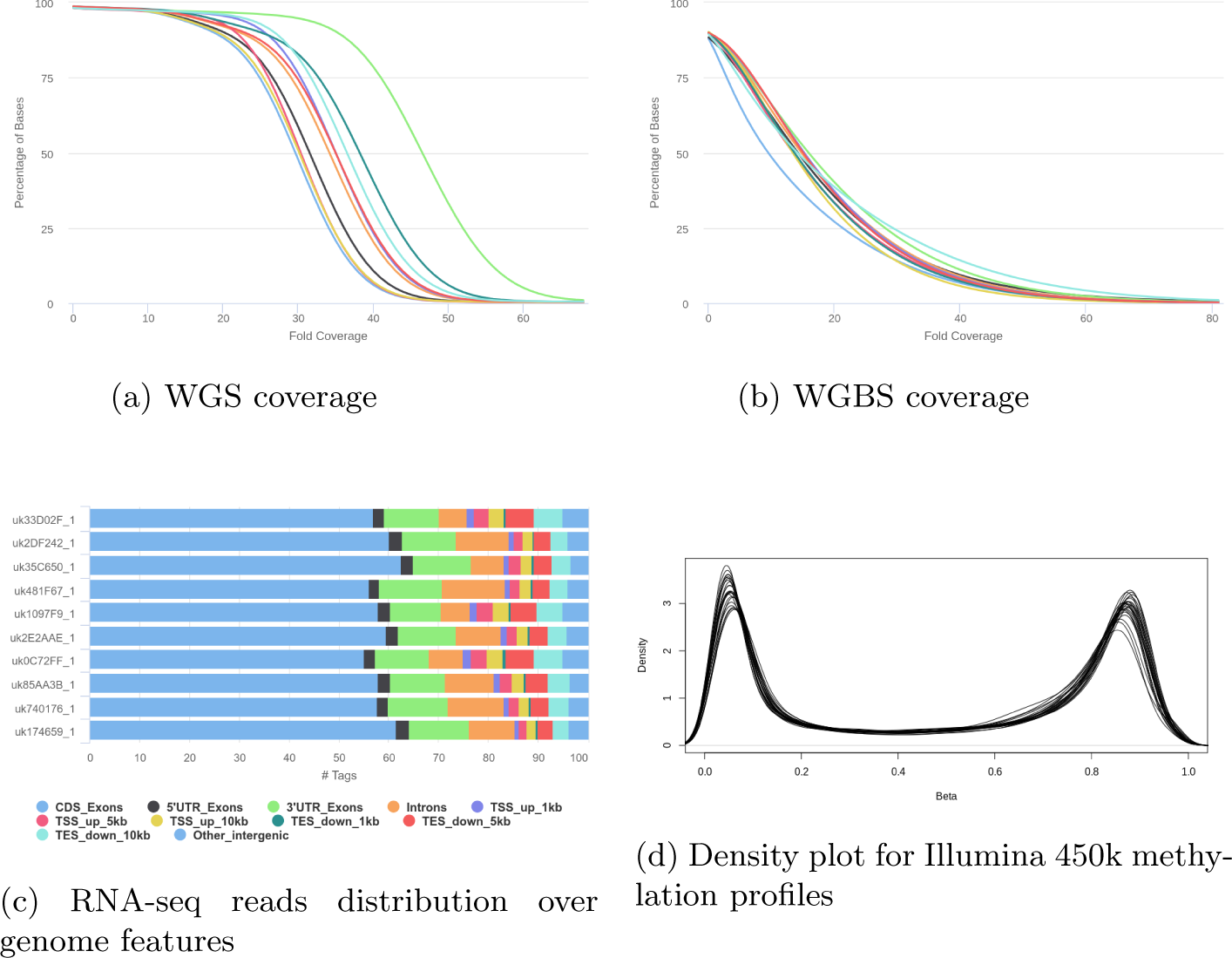
PGP-UK QC images for WGS, WGBS, RNA-seq and 450k methylation data

**Table 2:**
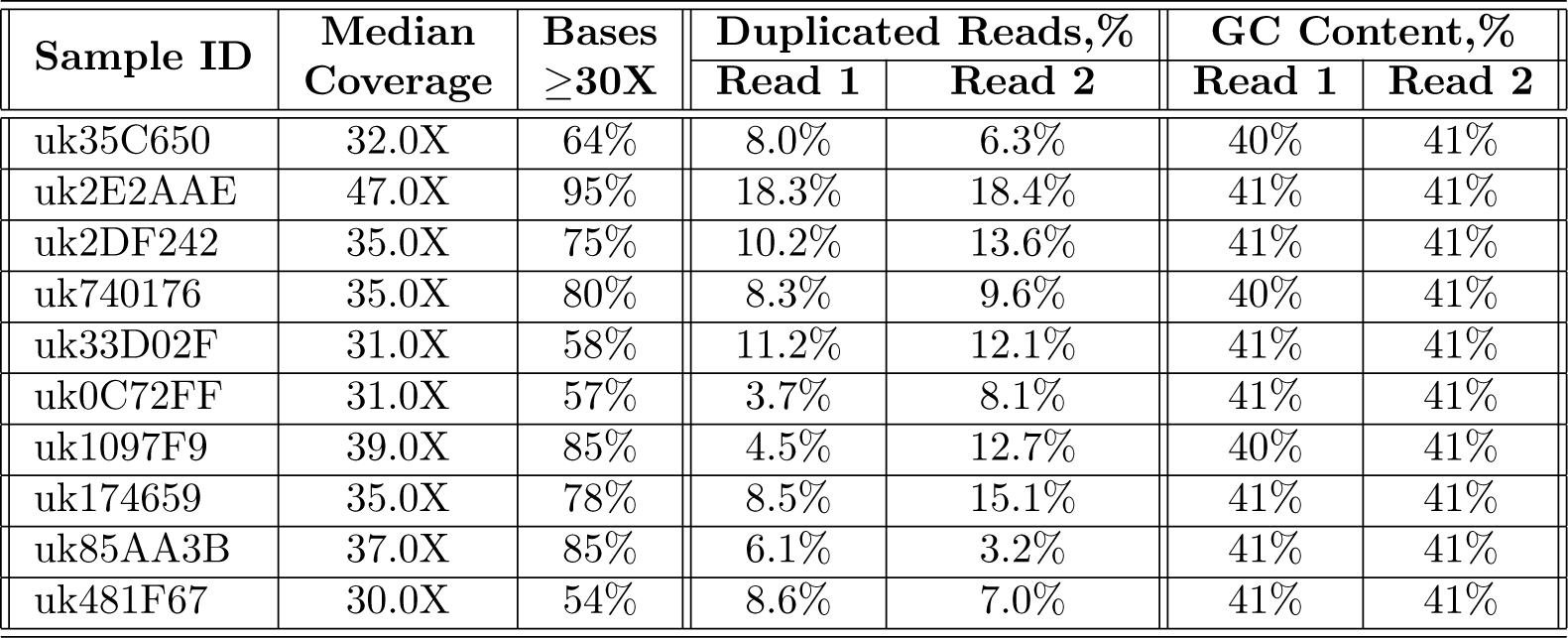
Quality control characteristics summary of the WGS data for PGP-UK participants.

| Sample ID | Median Coverage | Bases $\geq 30X$ | Duplicated Reads,% | | GC Content,% | |
| --- | --- | --- | --- | --- | --- | --- |
|  |  |  | Read 1 | Read 2 | Read 1 | Read 2 |
| uk35C650 | 32.0X | 64% | 8.0% | 6.3% | 40% | 41% |
| uk2E2AAE | 47.0X | 95% | 18.3% | 18.4% | 41% | 41% |
| uk2DF242 | 35.0X | 75% | 10.2% | 13.6% | 41% | 41% |
| uk740176 | 35.0X | 80% | 8.3% | 9.6% | 40% | 41% |
| uk33D02F | 31.0X | 58% | 11.2% | 12.1% | 41% | 41% |
| uk0C72FF | 31.0X | 57% | 3.7% | 8.1% | 41% | 41% |
| uk1097F9 | 39.0X | 85% | 4.5% | 12.7% | 40% | 41% |
| uk174659 | 35.0X | 78% | 8.5% | 15.1% | 41% | 41% |
| uk85AA3B | 37.0X | 85% | 6.1% | 3.2% | 41% | 41% |
| uk481F67 | 30.0X | 54% | 8.6% | 7.0% | 41% | 41% |

#### WGBS data QC

`GemBS` v.3.2.1, `FastQC` v.0.11.7 and `Picard` v.2.18.23 tools were used in quality control of the PGP-UK WGBS and data QC reports were generated using `MultiQC` v.1.5 software [8].

WGBS average median coverage is above 14X (ranging from 10X to 16X across samples) with more than 19% of bases covered reaching 30X or deeper (varies between 15% and 25% across samples), see Figure 2b. Summary of the WGBS QC analysis is presented in Table 3.

**Table 3:** Quality control characteristics summary of the WGBS data for PGP-UK participants.

| Sample ID | Median Coverage | Bases $\geq 30X$ | Duplicated Reads,% | | GC Content,% | |
| --- | --- | --- | --- | --- | --- | --- |
|  |  |  | Read 1 | Read 2 | Read 1 | Read 2 |
| uk35C650 | 10.0X | 15% | 27.3% | 13.3% | 26% | 29% |
| uk2E2AAE | 15.0X | 20% | 39.4% | 20.3% | 24% | 27% |
| uk2DF242 | 16.0X | 23% | 28.0% | 12.4% | 24% | 27% |
| uk740176 | 15.0X | 20% | 25.8% | 12.6% | 25% | 27% |
| uk33D02F | 16.0X | 20% | 26.3% | 13.1% | 24% | 27% |
| uk0C72FF | 14.0X | 18% | 26.8% | 11.4% | 25% | 28% |
| uk1097F9 | 14.0X | 15% | 26.0% | 10.8% | 24% | 27% |
| uk174659 | 14.0X | 17% | 27.1% | 15.5% | 24% | 27% |
| uk85AA3B | 16.0X | 19% | 28.3% | 14.9% | 24% | 27% |
| uk481F67 | 15.0X | 25% | 31.6% | 17.4% | 26% | 29% |

#### RNA-seq data QC

All of the RNA-seq samples were processed with a modified version of the `nextflow nf-core` RNA-seq pipeline (https://github.com/nf-core/rnaseq). Specifically, reads were trimmed with `TrimGalore` v.0.4.1, aligned against hg19 with `STAR` v.2.5.2a and duplicated reads were identified and removed with `Picard` v.2.18.9 tools. QC reports were generated using `MultiQC` v.1.5 [8] as part of the same pipeline. Reads were further split and trimmed using `GATK4`.

The mean RIN value of RNA used for sequencing was 8.55 (ranging between 7.1 and 9.3). Figure 2c demonstrates the distribution of mapped reads over various genomic features. A summary of the RNA-seq QC analysis is presented in Table 4.

**Table 4:** Quality control characteristics summary of the RNA-seq data for PGP-UK participants.

| Sample ID | RIN | Uniquely Aligned,% | Duplicated Reads,% |  | GC Content,% |  |
| --- | --- | --- | --- | --- | --- | --- |
|  |  |  | Read 1 | Read 2 | Read 1 | Read 2 |
| uk35C650 | 8.8 | 88.8% | 83.2% | 80.6% | 53% | 56% |
| uk2E2AAE | 9.1 | 89.3% | 85.9% | 82.3% | 53% | 56% |
| uk2DF242 | 9.2 | 90.0% | 86.3% | 81.9% | 53% | 56% |
| uk740176 | 8.5 | 90.0% | 84.8% | 80.6% | 53% | 56% |
| uk33D02F | 8.3 | 87.0% | 85.5% | 82.6% | 53% | 56% |
| uk0C72FF | 7.9 | 86.7% | 85.0% | 82.5% | 53% | 56% |
| uk1097F9 | 8.7 | 86.1% | 86.5% | 82.6% | 54% | 57% |
| uk174659 | 9.3 | 90.4% | 84.4% | 81.3% | 53% | 56% |
| uk85AA3B | 8.6 | 89.0% | 84.9% | 81.2% | 53% | 56% |
| uk481F67 | 7.1 | 90.4% | 87.3% | 83.7% | 52% | 55% |

#### 450k methylation data QC

450k DNA methylation profiles were generated from whole blood and saliva for each of the ten participants in the PGP-UK multi-omics reference panel. For quality control of these data, we used R v.3.5.2 with `minfi` v.1.28.3 and `ewastools` v.1.4 libraries [3, 9].

We performed quality checks based on 17 metrics assessed at control probes as described in the Illumina’s BeadArray Controls Reporter. All those checks were successfully passed. In addition, we analysed detection *p*-values and bead count information, which is available for 100% and 99.92% of CpGs respectively. 99.96% of the detection *p*-values are below the threshold of *α* = 0.01. Average CpG bead count number across all samples is 14, and 100% of the available bead count numbers ≥ 3. A summary of this analysis is presented in Table 5. Figure 2d shows the overlay of the *β*-value density distributions for all samples.

**Table 5:** Quality control characteristics summary (detection *p*-values and bead count) of the Illumina 450k data for PGP-UK participants.

| Sample ID | Tissue | Detection $p$-values | | Bead Count | |
| --- | --- | --- | --- | --- | --- |
| | | Available, % | $p < 0.01$, % | Available, % | $n \geq 3$, % |
| uk35C650 | blood | 100 | 99.98476 | 99.91370 | 100 |
|  | saliva | 100 | 99.97899 | 99.93710 | 100 |
| uk2E2AAE | blood | 100 | 99.92297 | 99.92503 | 100 |
|  | saliva | 100 | 99.93491 | 99.93670 | 100 |
| uk2DF242 | blood | 100 | 99.96478 | 99.90896 | 100 |
|  | saliva | 100 | 99.97178 | 99.92110 | 100 |
| uk740176 | blood | 100 | 99.91638 | 99.90361 | 100 |
|  | saliva | 100 | 99.91638 | 99.91800 | 100 |
| uk33D02F | blood | 100 | 99.92791 | 99.91761 | 100 |
|  | saliva | 100 | 99.92771 | 99.93120 | 100 |
| uk0C72FF | blood | 100 | 99.97467 | 99.90484 | 100 |
|  | saliva | 100 | 99.98558 | 99.88910 | 100 |
| uk1097F9 | blood | 100 | 99.98929 | 99.92460 | 100 |
|  | saliva | 100 | 99.98744 | 99.92150 | 100 |
| uk174659 | blood | 100 | 99.97714 | 99.94151 | 100 |
|  | saliva | 100 | 99.97899 | 99.92190 | 100 |
| uk85AA3B | blood | 100 | 99.93327 | 99.89351 | 100 |
|  | saliva | 100 | 99.94089 | 99.91670 | 100 |
| uk481F67 | blood | 100 | 99.98126 | 99.91514 | 100 |
|  | saliva | 100 | 99.98105 | 99.92130 | 100 |

### Multi-omics Data Matching

In order to ensure data integrity and exclude the possibility of sample mix-up between study participants, we validated our sample assignments, by matching the available 450k, WGBS and RNA-seq data against WGS. First, we matched the 450k against WGS data for each participant using 65 SNP control probes from Illumina 450k array. Second, we matched the WGBS-derived genotypes for the same 65 SNP loci with the WGS data. Third, we compared genotypes derived from RNA-seq and WGS data based on the set of loci from protein coding regions. The schema of the multi-omics data matching is given in Figure 3a and further details are provided below.

**Figure 3:**
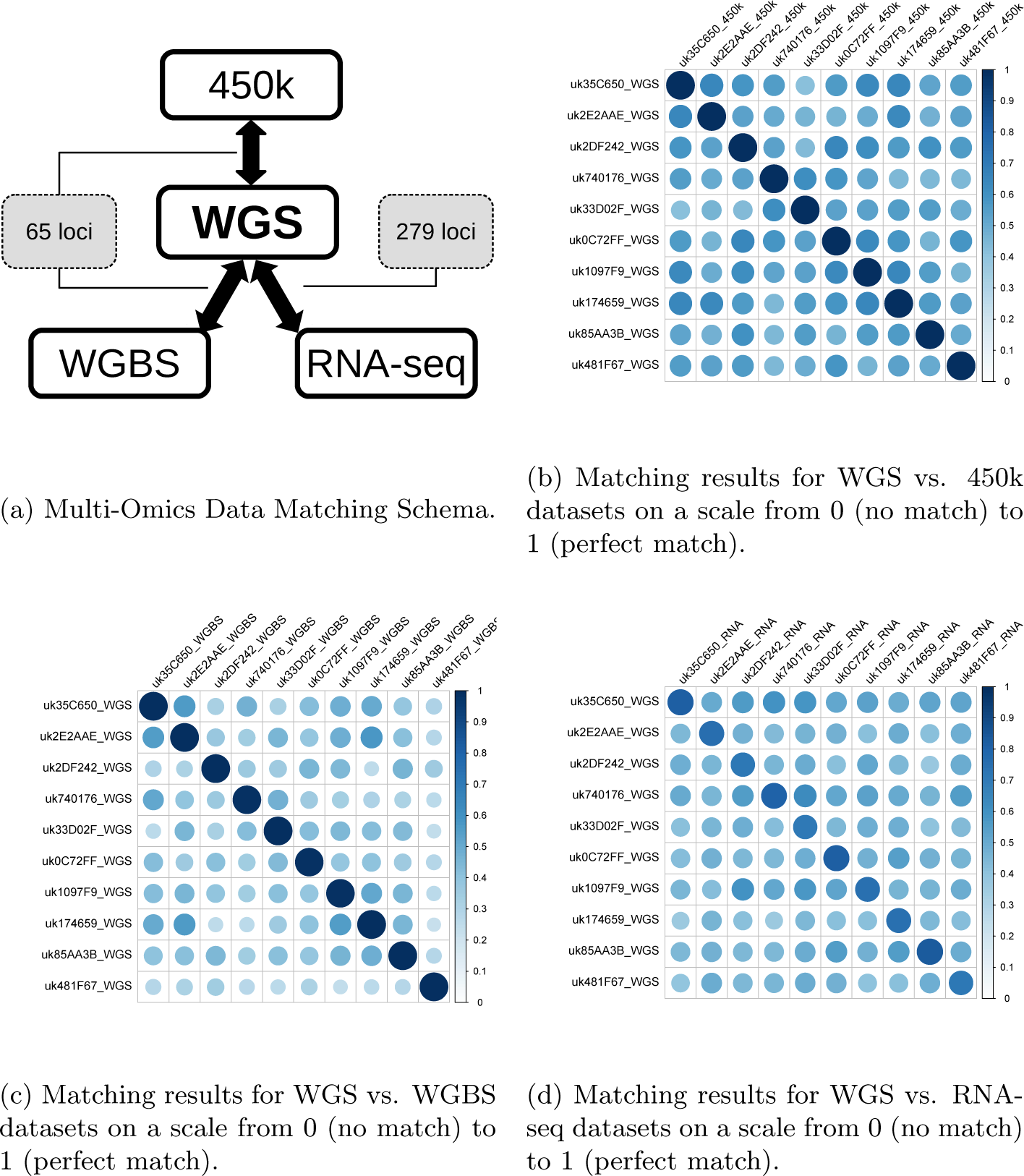
Multi-Omics Data Matching

We used *β*-values recorded at the 65 450k SNP control probes to distinguish between heterozygous and homozygous alleles in the 450k dataset. These SNPs are by design highly variable and can therefore provide a unique genetic signature that can be used to differentiate between each study participant. Note that 64 out of these 65 SNPs are outside protein-coding regions and, hence, not available for the RNA-seq data. Thus, we identified a list of 279 common loci (present in at least 4 WGS samples) that were located in the top 100 most expressed genes in the RNA-seq data, and that was used as validation set for the WGS vs. RNAseq comparison.

To match the different datasets, we extracted the locations of the loci used for validation (65 loci for the WGS vs. 450k and WGBS vs. 450k comparisons, and 279 loci for WGS vs. RNA-seq) and used the `HaplotypeCaller` and `GenotypeGVCFs` tools from (`GATK` v.3.8.0) on the corresponding BAM files to force the call of genotypes in these locations. Percentage of matching genotypes were then obtained across samples and datasets to confirm sample identity.

#### WGS vs. 450k

In order to obtain genotype information from 450k data, we extracted *β*-values for the 65 SNP control probes for each of the 10 PGP-UK participants. As expected, these *β*-values clustered into three separate peaks around 0.5 (which corresponds to heterozygous genotype), 0 and 1 (which correspond to homozygous genotype). We checked and confirmed that reported *β*-values for all SNP control probes which were derived from the whole blood and corresponding saliva 450k datasets were a 100% match. In other words, we established that the zygosity of each probe was the same across both DNA samples for any given participant.

We then extracted the genotypes for those 65 SNPs from WGS and matched them with to the corresponding zygosity in the 450k data. This resulted in perfect 100% match for corresponding samples, i.e. samples from the same participant.

#### WGS vs. WGBS

This comparison was performed by matching WGS- and WGBS-derived genotypes for 65 Illumina 450k array SNP control probes. The mean agreement between matched samples was 99.45%, which corresponds to a total of 3 loci mismatch observed in 3 out of ten participants (i.e. a single mismatch for each of those three participants). Altogether, 100% and 84.77% of 65 SNPs had coverage in the WGS and WGBS data respectively, which allowed us to make our comparison based on 51–61 common loci per participant.

#### WGS vs. RNA-seq

To match RNA-seq with WGS samples, we used a set of common loci in highly expressed genes as described above. Available genotypes for these loci were extracted from the RNA-seq and WGS samples and cross-validated. In total, 92.65% and 80.93% of these 279 loci had coverage in the RNA-seq and WGS data respectively, which allowed us to make our comparison based on 152-197 common loci per participant. On average, corresponding WGS and RNA-seq data are in agreement for 76.17% of genotype calls (range 69.68%–83.23%).

Results of matching 450k, RNA-seq and WGBS data with WGS are presented in Table 6. The correlation plots presented on Figure 3, demonstrate a substantially higher level of correspondence between samples from the same individual compared to those from different people when comparing WGS vs.450k (Figure 3b), WGS vs.WGBS (Figure 3c) and WGS vs.RNA-seq (Figure 3d).

**Table 6:** Summary of data cross-validation between 450k, WGBS and RNA-seq against WGS. Columns Loci, n and Loci, % contain respective numbers and percentages of loci used for matching (out of 65 loci for WGS and WGBS vs. 450k and 279 loci for WGS vs. RNA-seq).

| Sample ID | WGS vs. 450k |  |  | WGS vs.WGBS |  |  | WGS vs. RNA-seq |  |  |
| --- | --- | --- | --- | --- | --- | --- | --- | --- | --- |
|  | Loci, n | Loci, % | matched,% | Loci, n | Loci, % | matched,% | Loci, n | Loci, % | matched,% |
| uk35C650 | 65 | 100 | 100 | 52 | 80 | 100 | 161 | 57.71 | 81.99 |
| uk2E2AAE | 65 | 100 | 100 | 51 | 78.46 | 100 | 172 | 61.65 | 75.58 |
| uk2DF242 | 65 | 100 | 100 | 58 | 89.23 | 100 | 183 | 65.59 | 70.49 |
| uk740176 | 65 | 100 | 100 | 61 | 93.85 | 98.36 | 152 | 54.48 | 80.26 |
| uk33D02F | 65 | 100 | 100 | 53 | 81.54 | 100 | 188 | 67.38 | 69.68 |
| uk0C72FF | 65 | 100 | 100 | 57 | 87.69 | 98.25 | 159 | 56.99 | 81.13 |
| uk1097F9 | 65 | 100 | 100 | 54 | 83.08 | 98.15 | 190 | 68.10 | 73.68 |
| uk174659 | 65 | 100 | 100 | 52 | 80 | 100 | 197 | 70.61 | 74.62 |
| uk85AA3B | 65 | 100 | 100 | 60 | 92.30 | 100 | 167 | 59.86 | 83.23 |
| uk481F67 | 65 | 100 | 100 | 53 | 81.54 | 100 | 169 | 60.57 | 71.01 |

## Usage Notes

In this section, we describe two key outputs generated for each PGP-UK participants, the Genome and Methylome Reports. These reports are freely available to download on PGP-UK website, see https://www.personalgenomes.org.uk/data/.

Genome Reports leverage the information from variant call files (vcf) and provide an overview of the potential influence of genetic variants on several genetic traits, as well as ancestry information. Potentially beneficial or harmful traits for each participant were identified using public data from SNPedia [6], gnomAD v2.0.2 [12], GetEvidence [4] and ClinVar [11]. Plots to visualise the ancestry of each participant were created by applying principal component analysis (as implemented in `Plink` v1.9 [19]) on a genotype matrix resulting from merging the participant genotypes with those from 2504 unrelated samples from 26 worldwide populations available from the 1000 Genomes Project [1]. Population membership proportions were obtained using the `Admixture` v1.3.0 software [2] on above-mentioned genotype matrix.

Methylome reports contain epigenetic age and smoking status prediction for PGP-UK participants based on their methylome as assessed by 450k array experiments. Raw data (IDAT files) were processed, quality controlled and analysed using `ChAMP` [17, 20] and minfi [3] pipelines for `R`. Epigenetic age calculation was based on the multi-tissue Horvath clock [10], which predicts age using a linear combination of the methylation levels from a reference panel of 353 CpGs. Smoking status was predicted by calculating smoking scores as linear combinations of the methylation levels at 183 CpGs and then comparing them to a particular threshold as described in [7]. More details on the PGP-UK Genome and Methylome reports are described in [18].

## Acknowledgements

The authors acknowledge the use of the UCL Legion High Performance Computing Facility (Legion@UCL) and associated support services. We would like to thank UCL Genomics for array preparation and processing. PGP-UK gratefully acknowledges support from the Frances and Augustus Newman Foundation and the National Institute for Health Research (NIHR) UCLH Biomedical Research Centre (BRC369/CN/SB/101310). The views expressed are those of the author(s) and not necessarily those of the NIHR or the Department of Health and Social Care.

## Author contributions

OC wrote the manuscript with input from all authors. OC, AB, LC, JAGA, ELC, IM, RH, YT, VV and APW contributed analyses. JH and SB supervised the study. All authors read and approved the manuscript.

## Data Citations

1. *European Nucleotide Archive* PRJEB17529.
2. *ArrayExpress* E-MTAB-6523.
3. *European Nucleotide Archive* PRJEB25139.
4. *ArrayExpress* E-MTAB-5377.

